# Markonv: a novel convolutional layer with inter-positional correlations modeled

**DOI:** 10.1101/2022.06.09.495500

**Authors:** Jing-Yi Li, Yuhao Tan, Zheng-Yang Wen, Yu-Jian Kang, Yang Ding, Ge Gao

## Abstract

Deep neural networks equipped with convolutional neural layers have been widely used in omics data analysis. Though highly efficient in data-oriented feature detection, the classical convolutional layer is designed with inter-positional independent filters, hardly modeling inter-positional correlations in various biological data. Here, we proposed Markonv layer (Markov convolutional neural layer), a novel convolutional neural layer with Markov transition matrices as its filters, to model the intrinsic dependence in inputs as Markov processes. Extensive evaluations based on both synthetic and real-world data showed that Markonv-based networks could not only identify functional motifs with inter-positional correlations in large-scale omics sequence data effectively, but also decode complex electrical signals generated by Oxford Nanopore sequencing efficiently. Designed as a drop-in replacement of the classical convolutional layer, Markonv layers enable an effective and efficient identification for inter-positional correlations from various biological data of different modalities. All source codes of a PyTorch-based implementation are publicly available on GitHub for academic usage.

## 1 Background

In biology, motifs usually refer to recurring patterns associated with particular functions[1, 2, 3]. While site-independent model like position weight matrix (PWM) is a commonly used representation of biological motifs[4, 5, 6, 7], inter-positional correlations are pervasive in functional elements, like transcription factors binding sites[8, 9, 10, 11, 12, 10],RNA structure [13] as well as protein domains[14].On the other hand, while feature correlation has been successfully identified by previous studies via stacking multiple convolutional, LSTM, and Transformer layers[15, 16, 17, 18, 19], such strategy requires additional parameters and hardly represents an intuitive interpretation as that of classical convolutional layer.

Here, we proposed the Markonv layer (Markov convolutional neural layer), as a generalized form of convolutional neural layer, which models inter-positional correlations as Markov processes directly. Case studies based on both synthetic and real-world data demonstrated Markonv is effective and efficient in identifying motifs with intrinsic correlation for various scenarios, with a biology-meaningful, interpretable representation. Of note, Markonv layers can be a drop-in replacement of the convolutional layer in neural networks and can be adopted by existing networks with minimal effort. A PyTorch-based implementation of Markonv is publicly available on https://github.com/gao-lab/Markonv_figures.

## 2 Methods

In this section, we introduce the Markonv operator and the Markonv layer based on the Markonv operator. Both the Markonv operator and the Markonv layer are inspired by the convolutional layer, so we adopt the term ‘kernel’ to denote the mechanism of modeling the Markov process.

### Notation

We first define a Markonv convolution kernel as *K*, 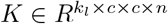. Where *k*_*l*_ represents the kernel length, *c* represents the number of channels (e.g., 4 for a one-hot encoded DNA sequence), and *n* represents the number of kernels. Then, we define the inputted sequence as *S, S* ∈ *R*^*b×c×l*^, Where *b* represents the batch size, and *l* represents the sequence length. Finally, the output obtained by the feedforward Markonv operator is referred to as *output, output*. 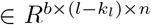 All coordinates are one-based (i.e., starting from one).

### Definition

For a given 0< *b*_0_ ≤ *b*, 0< *n*_0_ ≤ *n*, 0< *l*_0_ ≤ *l* − *k*_*l*_, the output is defined as:

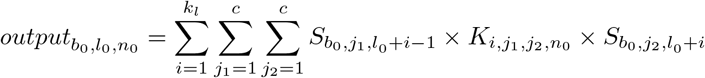

When using the chain rule to compute the gradients of the kernel and the inputted sequence, it can be proven that, for a given 0< *k*_0_ ≤ *k*, 0< *c*_0_ ≤ *c*, 0< *c*_1_ ≤ *c*, 0< *n*_0_ ≤ *n*, the gradient of *kernel* is:

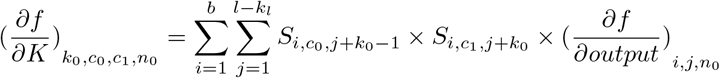

Where 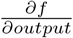 is the gradient of the output of Markonv.

Also, for a given 0< *b*_0_ ≤ *b*, 0< *c*_2_ ≤ *c*, 0< *l*_0_ ≤ *l*, the gradient of *input* is:

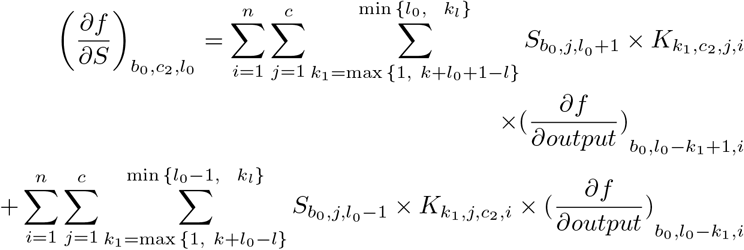

The Markonv kernel defined in this work fully determines the closed form solution of the (log-)probability of generating the sequence fragments from the corresponding first-order Markov process, given the observation of the initial state (see Appendix A for more details). We remove the probability of observing the initial state from the generation probability for simplicity. As a result, Markonv operator uses a new convolution kernel, which consists of a series of transition probability matrices, to compute the probability on the inputted sequence. As it scans the sequence for the first-order Markov process, it can extract inter-positional correlated features from the sequence.

We build the Markonv layer with the Markonv operator along with two optional modules for flexibility: the reverse sequence module and the boundary control module. The reverse sequence module reverses the inputted sequence to find the Markov process in the opposite direction. The boundary control module uses the same strategy as vConv[20] to adjust the kernel length adaptively during training (see Appendix B for more details).

## 3 Results

### 3.1 Benchmark datasets

In this section, we tested the performance of Markonv on three different datasets:

1. To test whether Markonv-based networks could model first-order Markov processes and converge with popular optimizers, we constructed four different simulation datasets.
2. To test whether Markonv-based network could be trained with less complexity than convolution-based networks on omics data while still obtaining a comparable performance, we compared Markonv-based networks with HOCNNLB[21] on RNA-binding protein (RBP) datasets.
3. To test whether Markonv-based network could be efficient in mining continuous input data, we compared our Markonv-based basecaller with Bonito (https://github.com/nanoporetech/bonito) on Oxford Nanopore datasets.

#### Simulation datasets

We constructed four different motifs, each of which is a Markov process (see Appendix C for more details). The Shannon entropy, a metric of motif chaoticness, gradually increases from the first motif to the fourth motif. We simulated for each motif case 6,000 sequences (with 3,000 positive and 3,000 negative) of length 1,000, picked 600 as the test dataset randomly, and randomly split the rest into 4,860 training and 540 validation sequences. The name of the dataset describes the index of the motif used; for example, Dataset “1” means that the dataset contains the first motif. Each negative sequence is a random sequence whose bases were independently sampled from the categorical distribution P(A)=P(C)=P(G)=P(T)=0.25. For each positive sequence, we first constructed a random sequence as described above, and then substituted a sequence fragment generated from one of the motifs associated with the case for a randomly chosen fragment of the same length.

#### RBP datasets

We downloaded the original dataset used by Zhang et al. from https://github.com/NWPU-903PR/HOCNNLB, including the training set and test set[21]. We randomly selected 20% of the training set to construct a validation set. At the same time, we downloaded the results of different neural network structures from Zhang et al. (https://ars.els-cdn.com/content/image/1-s2.0-S0003269719303513-mmc1.docx).

#### Oxford Nanopore datasets

We used the training and validation dataset in Bonito release v0.5.1 (https://github.com/nanoporetech/bonito). We used a benchmarking read set from *Klebsiella pneumoniae* as test set, as suggested by Wick et al.[22].

### 3.2 Network structures in each experiment

The networks we established and the baseline methods to compare within each experiment are elucidated below.

#### Networks for simulation datasets

We built a simple Markonv-based network with a Markonv layer, a global max-pooling layer, and a dense layer (see model structure details in Appendix D). The baseline method is the convolution-based network with the same structure.

#### Networks for RBP datasets

For Markonv-based networks, we followed the construction in the simulation case above. Baseline methods are HOCNNLB-1 to HOCNNLB-4, where “HOCNNLB-*k*” represents that the network encodes *k*-mer nucleotide as a one-hot vector and models the inputted sequence as a (*k* − 1)-order Markov process (if *k* ≥ 2), or just leaves the input as-is (if *k* is 1). These models are all neural networks mentioned in Zhang et al.’s work [21].

#### Networks for Oxford Nanopore datasets

Inspired by Bonito, we built a Markonv-based basecaller (Fig. 1). The baseline method is Bonito v3 network (config “dna_r9.4.1_e8_sup@v3.3” in Bonito release v0.5.1). The details of the training method are in Appendix E.

**Figure 1.**
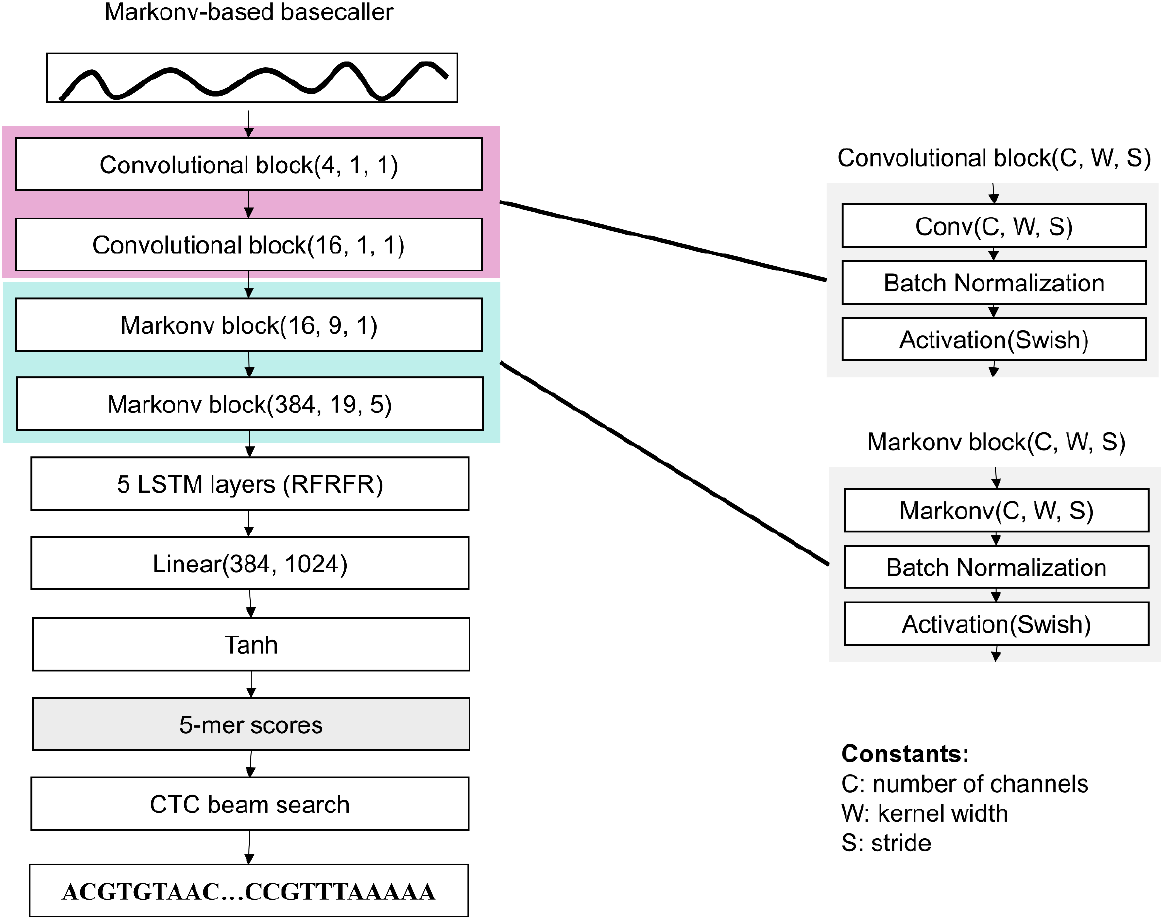
The architecture of the Markonv-based basecaller. We used the first two convolutional blocks to decode electric signals. “RFRFR” represents that there are five layers of LSTM, where “R” stands for reverse-LSTM, and “F” stands for forward-LSTM. 5-mer scores is the output of Tanh layer, which is a sequence with the values at each position describing the probability of each 5-mer at that position. We use CTC (Connectionist Temporal Classification) algorithm[23] to train the model; briefly, when training, we calculated the CTC loss using the 5-mer scores and the true sequences, and when testing, we used CTC beam search with a beam width 32 to decode sequences from 5-mer scores.

### 3.3 Results

#### Markonv-based networks outperformed convolution-based networks on simulation datasets

We compared the AUROC of the models on four simulation datasets generated by different motifs (Fig. 2). As the Shannon entropy of the motif increases, the amount of information contained in the motif decays rapidly, associated with the observation that the AUROC of the Markonv-based network becomes higher than that of the convolution-based network. When the data inserted in the dataset was relatively simple (Dataset 1), the Markonv-based network had a performance similar to that of the convolution-based network, but as the complexity of the motif increased (Datasets 2, 3, and 4), the Markonv-based network was statistically better than the convolution-based network, with the *p*-value of the two-tailed Wilcoxon’s Rank Sum test less than 0.001.

**Figure 2.**
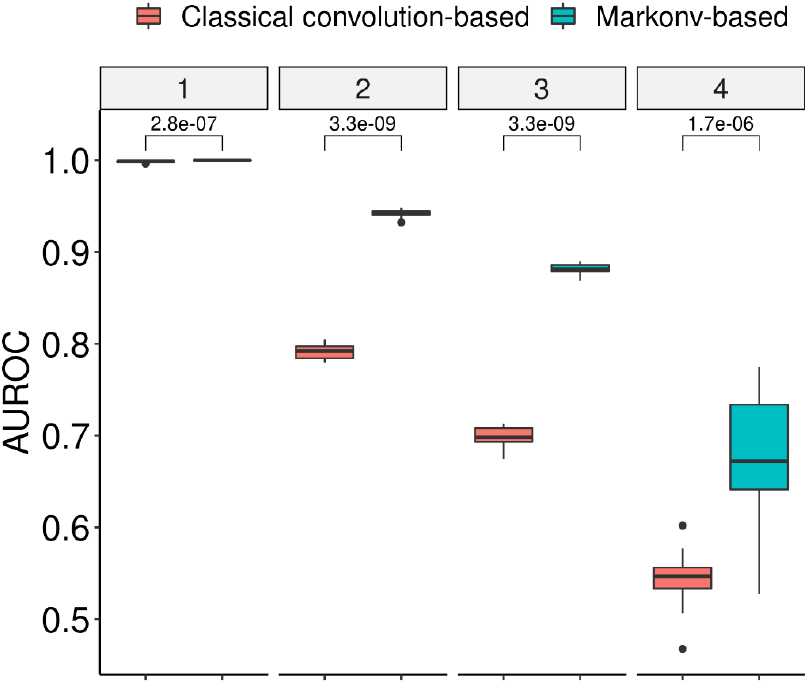
Comparison of Markonv-based networks and convolution-based networks on simulation datasets. The numbers in the facet labels represent the dataset indices (see Section 3.1 for more details). The boxplots under each facet describe the distribution of AUROCs of the networks trained with different random seeds on the same dataset. *P*-values shown above each pair of boxplots were computed by two-tailed Wilcoxon’s Rank Sum test and unadjusted.

We selected for each Markonv-based network those Markonv kernels that are predictive (see Appendix F for details of the method), and recovered a first-order Markov process similar to the inserted real motif (Fig. 3; see Appendix G for the comparison between all pairs of real and recovered motifs).

**Figure 3.**
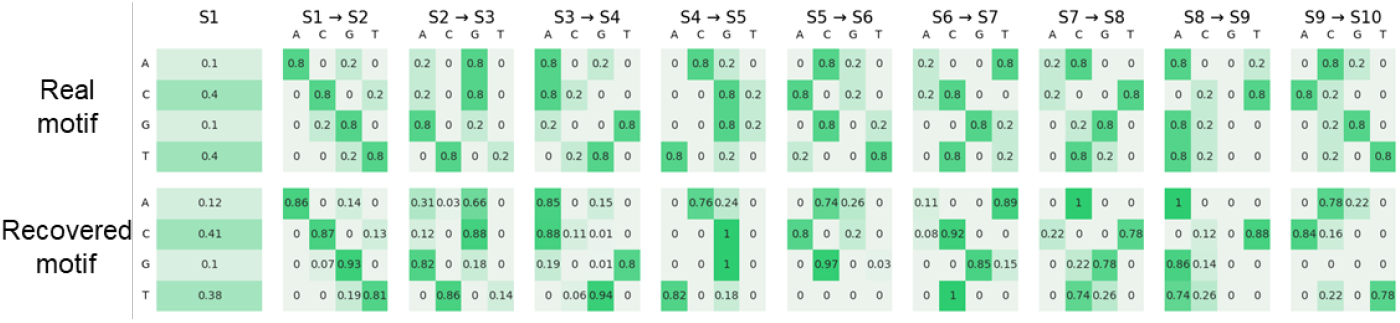
Motifs recovered using Markonv kernels on Datasets 3. The “Real motif” represents the motif inserted into the inputted sequences, and the “Recovered motif” represents the motif recovered from the kernels. “S1” represents the probability of observing each base at the first position, and “Si → S(i+1)” represents the Markov transition matrix from the i-th position to the (i+1)-th position, with the element at j-th row and k-th column being the probability of observing the base described by the k-th column name at the (i+1)-th position under the condition that the base described by the j-th row name is observed at the i-th position.

#### Markonv-based networks could use fewer parameters to outperform the more complex HOCNNLB models on classifying RBP datasets

Fig.4 displays the distribution of different networks’ AUROC across all RBP datasets. While the Markonv-based network has lower space complexity than HOCNNLB models (Table 1), it not only outperformed the simple convolutional neural network (HOCNNLB-1), but also the one designed to model the first-order Markov process (HOCNNLB-2), and even had a performance close to those modeling higher-order representation (HOCNNLB-3 and HOCNNLB-4)

**Figure 4.**
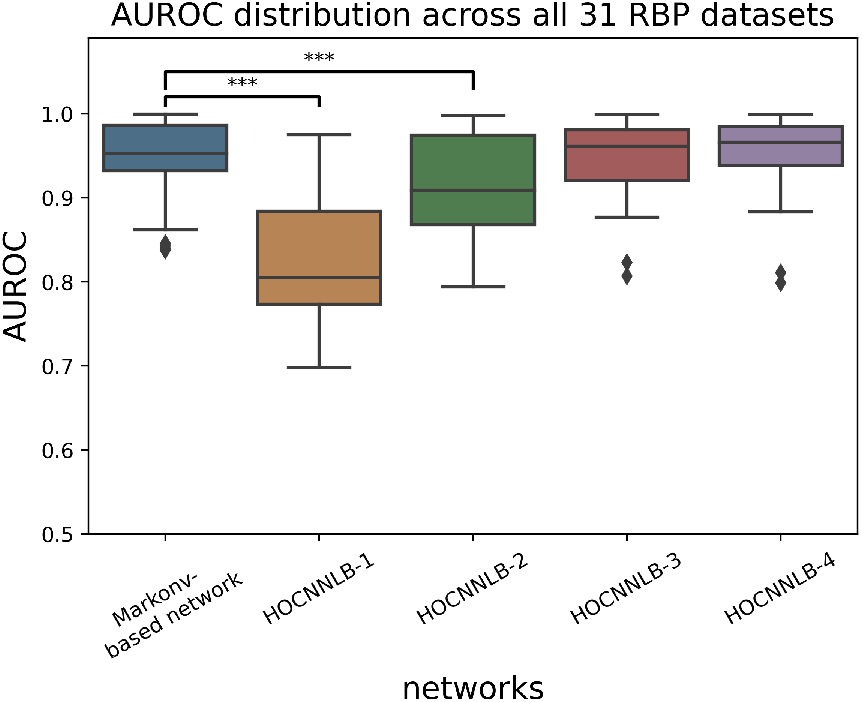
Comparison of AUROC between the Markonv-based network and other networks. Boxplots represent the AUROC distribution of each network across all 31 datasets. Details of each network are available in Section 3.2. ***, an unadjusted *p*-value of two-tailed Wilcoxon’s Rank Sum test that is less than 0.001.

**Table 1.** Comparison of the computational complexity of different neural layers. where *N* represents the number of kernels, *l* represents the length of the convolution kernel, *n* represents the length of the inputted sequence, *c* represents the number of channels, and *k* represents the order of HOCNNLB.

| Name | Computational complexity | Space complexity |
| --- | --- | --- |
| Convolutional layer | $O(N * l * c * n)$ | $O(N * l * c * n)$ |
| Markonv | $O(N * l * c^2 * n)$ | $O(N * l * c * n)$ |
| HOCNNLB-$k$'s convolutional layer | $O(N * l * c^k * n)$ | $O(N * l * c^k * n)$ |

#### Markonv-based basecaller rivaled Bonito’s basecaller performance with much fewer parameters

Table 2 summarizes the performance of the Bonito and Markonv-based basecaller. Although trained with the same default optimizer and learning rate scheduler, the Markonv-based basecaller could still use much fewer parameters than Bonito to achieve a comparable performance, suggesting Markonv’s potential broad application to various types of biological data.

**Table 2.** Performance of the Bonito and Markonv-based basecaller. Read accuracy is the sequence identity between each reference read and its basecalled result; similarly, consensus accuracy is the sequence identity between the reference and the consensus sequence (which is assembled from overlapping basecalled results).

| Name | Median read accuracy | Median consensus accuracy | Number of network parameters |
| --- | --- | --- | --- |
| Bonito | 92.12% | 99.92% | 27,008,104 |
| Markonv-based basecaller | 92.44% | 99.94% | 9,115,728 |

## 4 Discussion

As an attempt to identify complex motifs with intrinsic correlation effectively and efficiently, we here proposed Markonv, a novel convolutional layer combining the classical convolution operator and the Markov generation process. Experiments on simulation datasets demonstrated that Markonv layers could recover first-order Markov processes and improve the model performance. The evaluation on RBP datasets and Oxford Nanopore sequencing datasets demonstrated that Markonv-based networks achieved a comparable performance with fewer parameters than classical convolution-based networks, suggesting its potential advantage in reducing model complexity. The loss curve of each model on the simulation dataset shows that the convergence speed of the Markonv-based network was comparable to that of the classical convolution-based network, suggesting Markonv did not increase the optimization difficulty (Fig. 5; see Appendix H for details of loss curve for each dataset).

**Figure 5.**
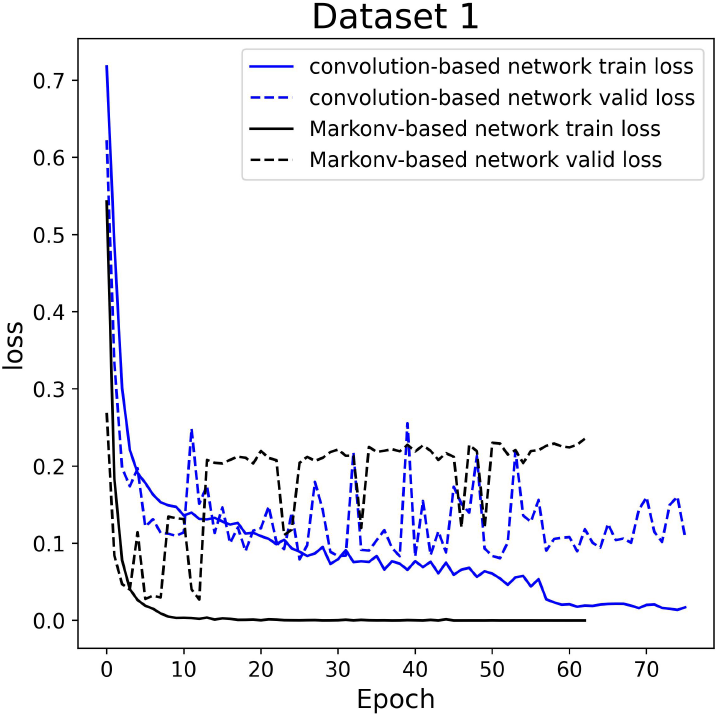
The training and the validation loss curves (with respect to the number of epochs) of the Markonv-based network and the convolution-based network on simulation Dataset 1.

Markonv is a neural layer that could be used as a module for processing one-dimensional sequence data to replace the classical convolutional neural layer or (in theory) the recurrent neural layer in the neural network. Compared to convolutional neural layers, Markonv can model first-order Markov processes with only one layer. Compared with the parameters in recurrent neural layers, the Markonv kernel is interpretable in the sense that an equivalent Markonv process can be constructed from the kernel (see Appendix F-G) for subsequent data analysis. Our work has opened some meaningful directions to be further exploited: (i) although we have provided the loss curves to demonstrate the speed of convergence for Markonv, the theoretical guarantee of convergence (if exists) is still needed; (ii) the results on RBP datasets suggested that biological sequences contain higher-order representations, so it would be meaningful to expand the idea of Markonv to model higher-order representations.

## Notes

### Competing Interest Statement

The authors have declared no competing interest.

