## supplementary files for "Markonv: a novel convolutional layer with inter-positional correlations modeled"

#### A Theorem of Markonv operator

**Theorem1.** The Markonv convolution kernel-$K \in R^{k_l \times c \times c \times n}$ fully determines the closed form solution of the (log-)probability of generating the sequence fragments from the corresponding first-order Markov process, given the observation of the initial state.

*Proof.*

Given the fact that the convolution processes of different sequences and different kernels are independent, and that the convolution processes of the same kernel on sequences are independent of each other, it suffices to prove that for any given kernel and any given sequence fragment, the value under the Markonv operator of the fragment and the kernel is equal to the linear transformation of the log-probability value of the sequence generated by a particular Markov process, where this Markov process is determined by the given kernel.

Therefore, we only focus on a single kernel K: $= K_{:,:,:,1} \in R^{k_l \times c \times c}$ , and the single sequence fragment F $:= S_{1,:,[1,\ldots,(k_l+1)]} \in R^{c \times (k_l+1)}$, where F is one-hot encoded (i.e., $\forall 1 \leq i \leq k_l + 1$ and $1 \leq j \leq c$ , we have $\mathrm{F}_{j,i} \in \{0,1\}$ and $\sum_{h=1}^{c} \mathrm{F}_{h,i} = 1$). We use the index sequence $\{J(1), J(2), \ldots, J(k_l+1)\}$ to describe the one-element positions in F (i.e., they satisfy $\mathrm{F}_{J(i),i} = 1$).

The corresponding first-order Markov process we'd like to relate to this kernel is then defined by the following series of transition matrices $f(\mathrm{K}, b) \in R^{k_l \times c \times c}$ , where $b > 1$ and $f(\mathrm{K}, b)_{i,j_1,j_2} := \frac{b^{(\mathrm{K})_{i,j_1,j_2}}}{\sum_{j'_2=1}^{c} b^{(\mathrm{K})_{i,j_1,j'_2}}}$. It is thus clear that each matrix $f(\mathrm{K}, b)_{i,:,:}$ is indeed a transition matrix related to the position $i$, because for all possible combinations of $i, j_1, j_2$, we always have (1) $f(\mathrm{K}, b)_{i,j_1,j_2}$ lies within $[0, 1]$ and (2) $\sum_{j_2=1}^{c} f(\mathrm{K}, b)_{i,j_1,j_2} = 1$.

Then the closed form solution of the log-probability of generating the sequence fragment (F) from the corresponding first-order Markov process (defined by $f(\mathrm{K}, b)$ ), given the observation of the initial state (i.e., we do not compute the probability for this observation), will be:

$$
\begin{aligned}
&\ln\left(P\left(\mathrm{F} \mid f\left(\mathrm{K}, b\right)\right)\right)\\
&= \ln\left(\prod_{i=1}^{k_l}\prod_{j_1=1}^{c}\prod_{j_2=1}^{c}\left(\left(\mathbf{1}_{\mathrm{F}_{j_1,i}=1} * \mathrm{F}_{j_1,i}\right) * \left(\mathbf{1}_{\mathrm{F}_{j_2,i+1}=1} * \mathrm{F}_{j_2,i+1}\right) * f(\mathrm{F} \mid f(\mathrm{K}, b))\right)\right)\\
&= \ln\left(\prod_{i=1}^{k_l}\left(\mathrm{F}_{J(i),i} * \mathrm{F}_{J(i+1),i+1} * f(\mathrm{K}, b)_{i,J(i),J(i+1)}\right)\right)\Bigg)\\
&= \ln\left(\prod_{i=1}^{k_l} f(\mathrm{K}, b)_{i,J(i),J(i+1)}\right)\\
&= \sum_{i=1}^{k_l}\ln\left(\frac{b^{(\mathrm{K})_{i,J(i),J(i+1)}}}{\sum_{j'_2=1}^{c} b^{(\mathrm{K})_{i,j(i),j'_2}}}\right)\\
&= (\ln(b))\sum_{i=1}^{k_l}(\mathrm{K})_{i,J(i),J(i+1)} - \sum_{i=1}^{k_l}\ln(\sum_{j'_2=1}^{c} b^{(\mathrm{K})_{i,J(i),j'_2}}).
\end{aligned}
$$

On the other hand, the output, K ■ F, is the following scalar:

$$
\mathrm{K} \blacksquare \mathrm{F} = \sum_{i=1}^{k_l}\sum_{j_1=1}^{c}\sum_{j_2=c}^{c} S_{1,j_1,i} * K_{i,j_1,j_2,1} * S_{1,j_2,i+1}
$$

(By definition of Markonv operator)

$$
= \sum_{i=1}^{k_l} \mathrm{F}_{J(i),i} * \mathrm{K}_{i,J(i),J(i+1)} * \mathrm{F}_{J(i+1),i+1}
$$

(The other elements in F are 0 )

$$
= \sum_{i=1}^{k_l} \mathrm{K}_{i,J(i),J(i+1)}
$$

We then have:

$$\ln\left(P\left(\mathrm{F}\mid f\left(\mathrm{K},b\right)\right)\right) = \left(\ln\left(b\right)\right)\sum_{i=1}^{k_l}\left(\mathrm{K}\right)_{i,J(i),J(i+1)} - \sum_{i=1}^{k_l}\ln\left(\sum_{j'_2=1}^{c} b^{(\mathrm{K})_{i,J(i),j'_2}}\right)$$

$$= \left(\ln\left(b\right)\right)(\mathrm{K}\blacksquare\mathrm{F}) + d(\mathrm{K},b)$$

where $d(\mathrm{K},b) := -\sum_{i=1}^{k_l}\ln(\sum_{j'_2=1}^{c} b^{(\mathrm{K})_{i,J(i),j'_2}})$ fully depends on $\mathrm{K}$ and $b$ given the inputted sequence (the $J(i)$ values).

#### B The details of two optional modules for the Markonv layer

The boundary control module helps to adaptively learn the variable kernel length during training as vConv does[Li et al., 2021]; this is achieved by introducing boundary parameters, $w_m^{z,0}$ and $w_m^{z,1}$, to each of the Markonv kernels.(Fig. 1A). Specifically, assuming the length (along the positional axis) of the raw Markonv kernel is $L$ and the number of channels is 4, we generate the masked kernel as follows: (1) we first generate a $M \in R^{L\times4\times4}$ with $M_{i,j,l} = \left(1+e^{-(i-1-w_m^{z,0})}\right)^{-1}$, $\forall\, 0 < i \leq L,\ 0 < j \leq 4, 0 < l \leq 4$ ; the final $M_{i,j,l}$ is close to 0 when $i \gg w_m^{z,0}$; (2) similarly, we generate a $M^* \in R^{L\times4\times4}$ with ${M^*}_{i,j,l} = \left(1+e^{-(w_m^{z,1}-(i-1))}\right)^{-1}$, $\forall\, 0 < i \leq L,\ 0 < j \leq 4, 0 < l \leq 4$; the final ${M^*}_{i,j,l}$ is close to 0 when $i \ll w_m^{z,1}$ ; (3) we then compute the mask matrix (Fig. 1A) by taking $M + M^* - 1$ ; and (4) finally, we generate the masked kernel by masking this mask matrix onto the raw kernel, effectively masking the out-of-boundary elements to zeroes (Fig. 1A).

The reverse sequence module helps to identify Markov processes in the reverse direction of the inputted sequence (Fig. 1B); this is essential if the directionality of the underlying Markov processes is not guaranteed to follow the direction of the inputted sequence. In practice, this is achieved by applying a separate Markonv convolution kernel onto the original (forward) and the flipped (reverse) inputted sequences, and concatenating the result along the axis of the kernel index (Fig. 1B).

#### C The details of simulation motifs

We constructed four different motifs, each of which is a Markov process (Fig. 2). Each line represents a motif, and the chaoticness of the motif described by its (Shannon) entropy, increases from top to bottom. "S1" represents the probability of observing each base at the first position, and "Si $\rightarrow$ S(i+1)" represents the Markov transition matrix from the i-th position to the (i+1)-th position, with the element at j-th row and k-th column being the probability of observing the base described by the k-th column name at the (i+1)-th position under the condition that the base described by the j-th row name is observed at the i-th position.

#### D The structure of Markonv-based network

The structure of Markonv-based network we used in the manuscript (except for the Markonv-based basecaller) is shown in Fig. 3, consisting of a Markonv layer, a global max-pooling layer, a fully connected layer, and a sigmoid layer.

#### E Training method

**Training method for experiments on simulation dataset.** We used RMSprop[Hinton et al., 2012] for the optimization with learning rate 0.01. Both the convolutional neural layer and the Markonv layer in the model used the Glorot uniform initializer[Glorot and Bengio, 2010] to initialize the kernel. We used an early stopping mechanism (with patience being 50 epochs) when updating the parameters. The number of kernel is 128, the kernel length is 10, and each experiment was initialized with 16 different sets of parameters.

**Training method for experiments on RBP dataset.** We used RMSprop[Hinton et al., 2012] for the optimization with learning rate 0.001. We used an early stopping mechanism (with patience being

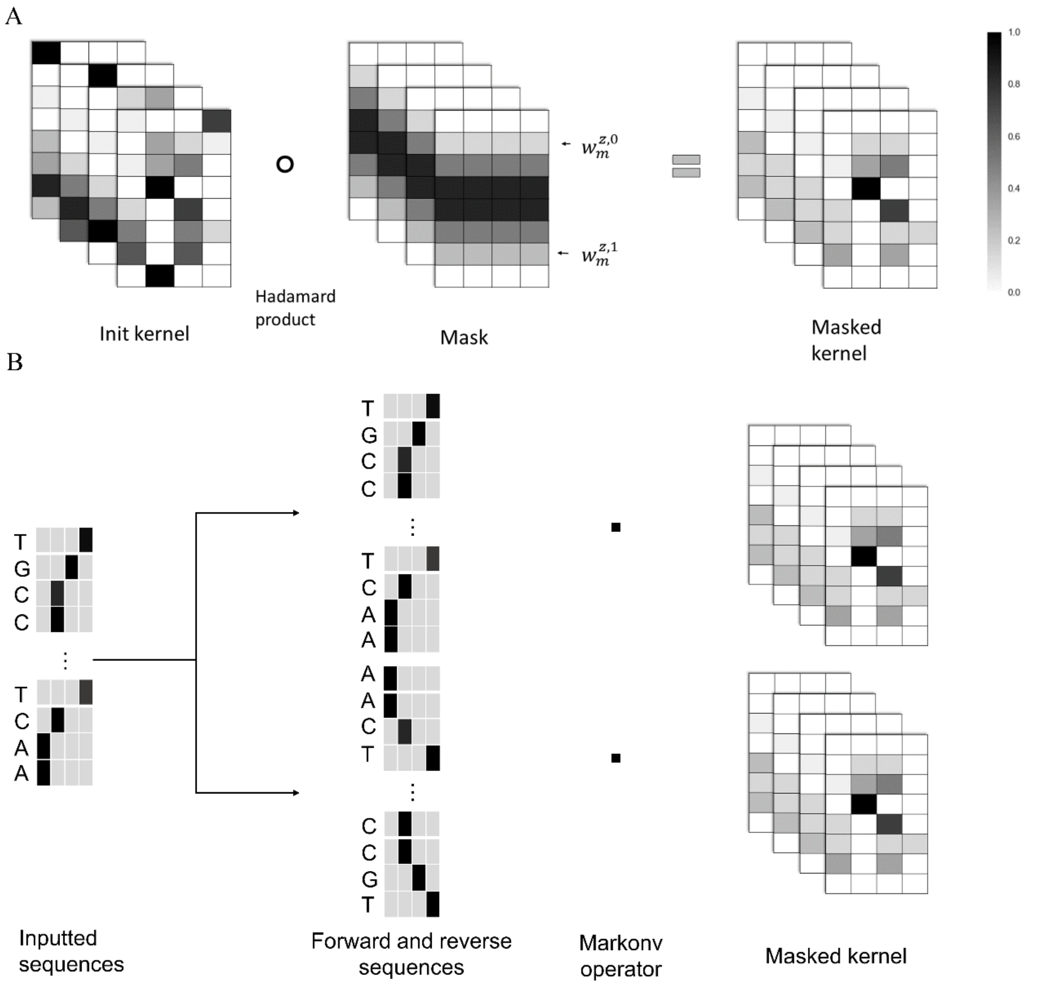


Figure 1: The two optional modules for the Markonv layer. (A) shows how the boundary control module uses the masked kernel to control the variable kernel length; (B) shows how the reverse sequence module helps to identify Markov processes in the reverse direction of the inputted sequence.

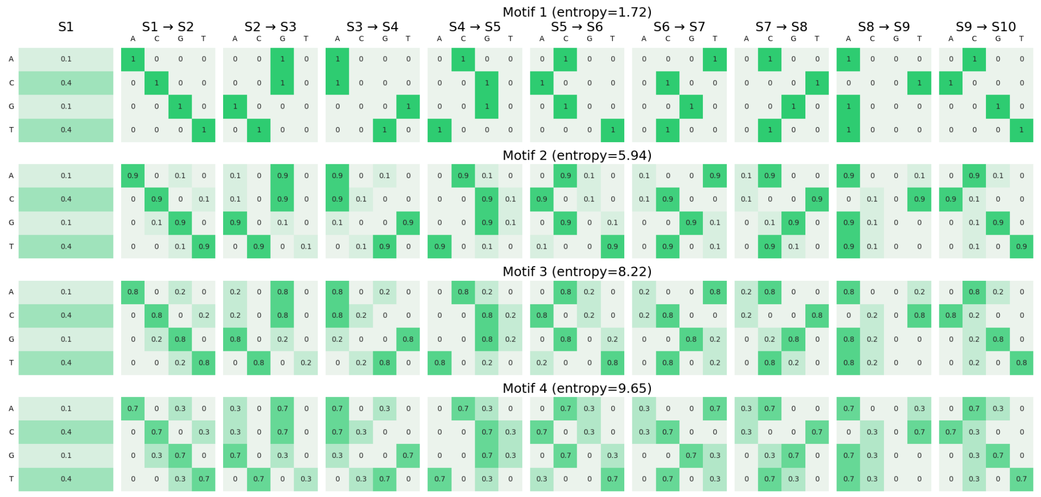


Figure 2: The Markov motifs used by the simulation dataset.

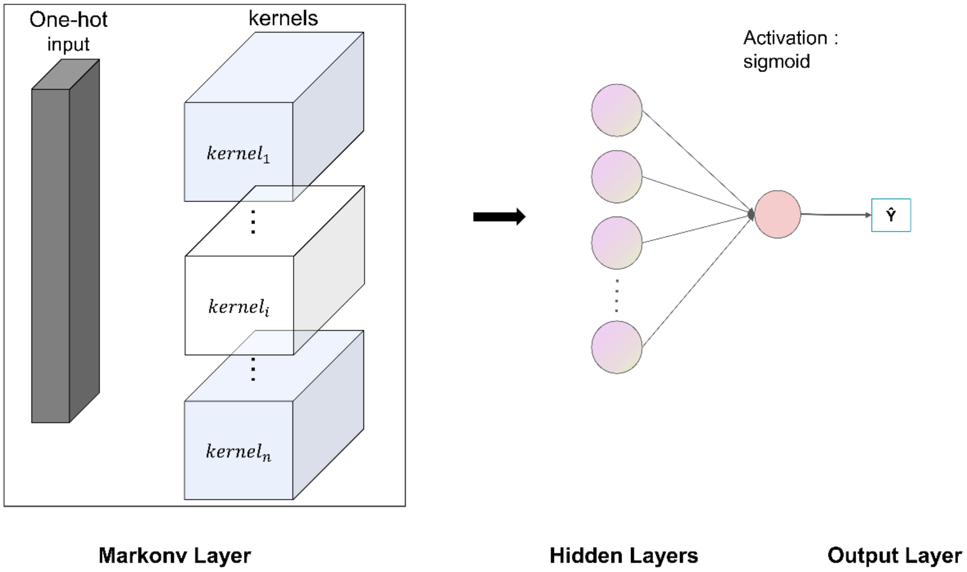


Figure 3: Structure of the (single-layer) Markonv-based network.

20 epochs) when updating the parameters. The number of kernel is 128, the kernel length is from {16,20}, and each experiment was initialized with 8 different sets of parameters.

**Training method for basecalling.** For Bonito and the Markonv-based network, we used the default hyperparameter settings in Bonito while setting training epochs to 15. Specifically, we used Adam optimizer [Kingma and Ba, 2014] with weight decay 0.01. We varied the learning rate according to the formula

$$lr = init\_lr * \begin{cases} 0.1 + 0.9 * \frac{step}{warmup\_steps}, & step < warmup\_steps \\ 0.01 + 0.5 * 0.99 * \left(\cos\left(\pi * \frac{step - warmup\_steps}{total\_steps - warmup\_steps}\right) + 1\right), & step \geq warmup\_steps \end{cases}$$

It corresponds to a linear warmup and a cosine decay of the learning rate. We used $init\_lr = 2 \times 10^{-3}$, and $warmup\_steps = 4000$. We trained the networks for 15 epochs and chose the parameters with the best performance on the validation set from these epochs.

**Computing resources and assets license.** We used some datasets in this paper, and we listed their copyrights as follows: the copyright of RBP datasets is Institute of Control & Information (ICI), Northwestern Polytechnical University, China Key Laboratory of Information Fusion Technology (LIFT), Ministry of Education, China; the copyright of Bonito training dataset is the Oxford Nanopore Technologies; the license of Bonito benchmark dataset is Creative Commons Attribution 4.0 (CC BY 4.0). All data is publicly available, and it neither has consent-related issues, nor contains personally identifiable information or offensive content. We referred to the code of some work in this paper, and we listed them as follows: the implementation of Bonito (`https://github.com/nanoporetech/bonito`) is distributed under the terms of the Oxford Nanopore Technologies, Ltd. Public License; the implementation of basecalling comparison (`https://github.com/rrwick/Basecalling-comparison`)[Wick et al., 2019] is made available under GNU General Public License. Experiments were conducted using RTX 3090 GPUs. It took several hours to finish experiments on simulation datasets and several days to finish experiments on Oxford Nanopore datasets with one RTX 3090.

### F Method for recovering motifs

We used a method similar to the classical motif recovering method for convolutional neural networks (Alipanahi 2015) (Fig. 4). In the first step, we selected the ten most important kernels in Markonv layer, where the importance for each kernel is defined as its own predictive performance of the

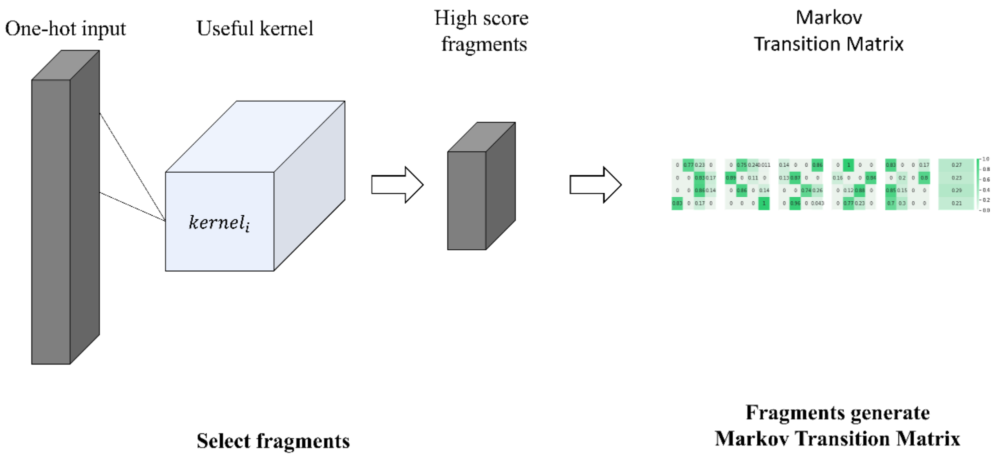


Figure 4: How to recover the transition matrix of the first-order Markov process from Markonv layer.

network with the outputs of all other kernels fixed to 0. In the second step, we extracted for each pair of (kernel, inputted sequence) the segment from the inputted sequence with the largest Markonv output value (called "generated score"). In the third step, we selected for each kernel those of its fragments whose generated scores are above the average of all its fragments' generated scores. Finally, we used the selected fragments of each kernel to generate transition matrices of the first-order Markov process for that kernel.

#### G Recovered motifs in simulation datasets

The recovered motifs in all simulation datasets are shown in Fig. 5. The "Real motif" represents the motif inserted into the input sequences, and the "Recovered motif" represents the motif recovered from the kernels. We only reported the recovered motif with the least difference from the real motif (measured by the Frobenius norm) across all kernels. The visualization of motifs follows that in Appendix C. For simplicity, we only visualized the part of the recovered motif that could be aligned to the real motif.

#### H Loss curve on each experiment

We recorded the loss curve of the Markonv-based network on each simulated dataset and the Markonv-based basecaller on Oxford Nanopore sequencing data. The results show that the optimization curves of Markonv-based network and the Markonv-based basecaller did not diverge much from those of the classical convolution-based network (for simulation datasets) or the Bonito model (for the Oxford Nanopore sequencing data) (Fig. 6).

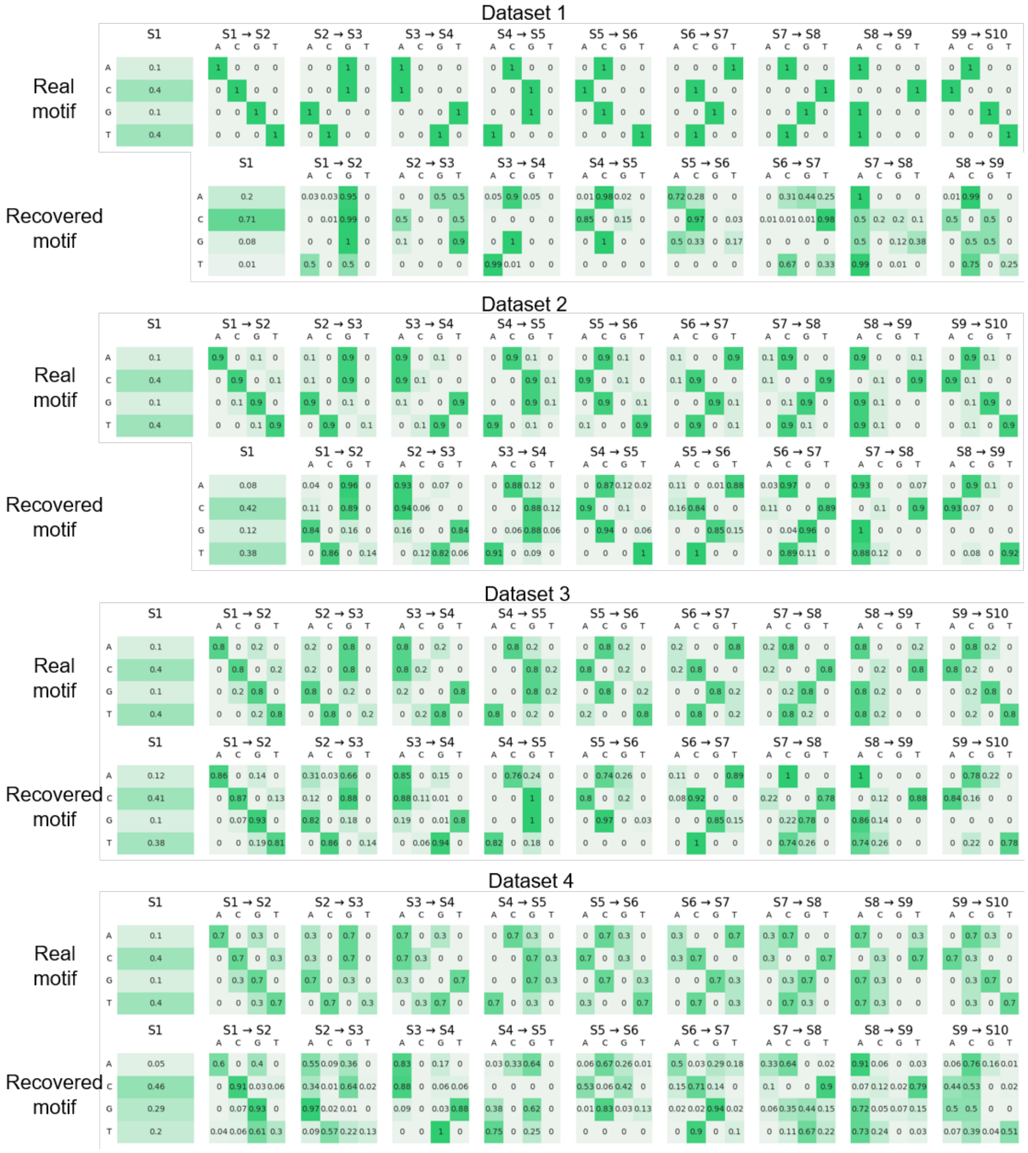


Figure 5: Motifs recovered by kernels on all simulation datasets.

Xavier Glorot and Yoshua Bengio. Understanding the difficulty of training deep feedforward neural networks. In *Proceedings of the thirteenth international conference on artificial intelligence and statistics*, pages 249–256. JMLR Workshop and Conference Proceedings, 2010.

Diederik P Kingma and Jimmy Ba. Adam: A method for stochastic optimization. *arXiv preprint arXiv:1412.6980*, 2014.

Ryan R Wick, Louise M Judd, and Kathryn E Holt. Performance of neural network basecalling tools for oxford nanopore sequencing. *Genome biology*, 20(1):1–10, 2019.

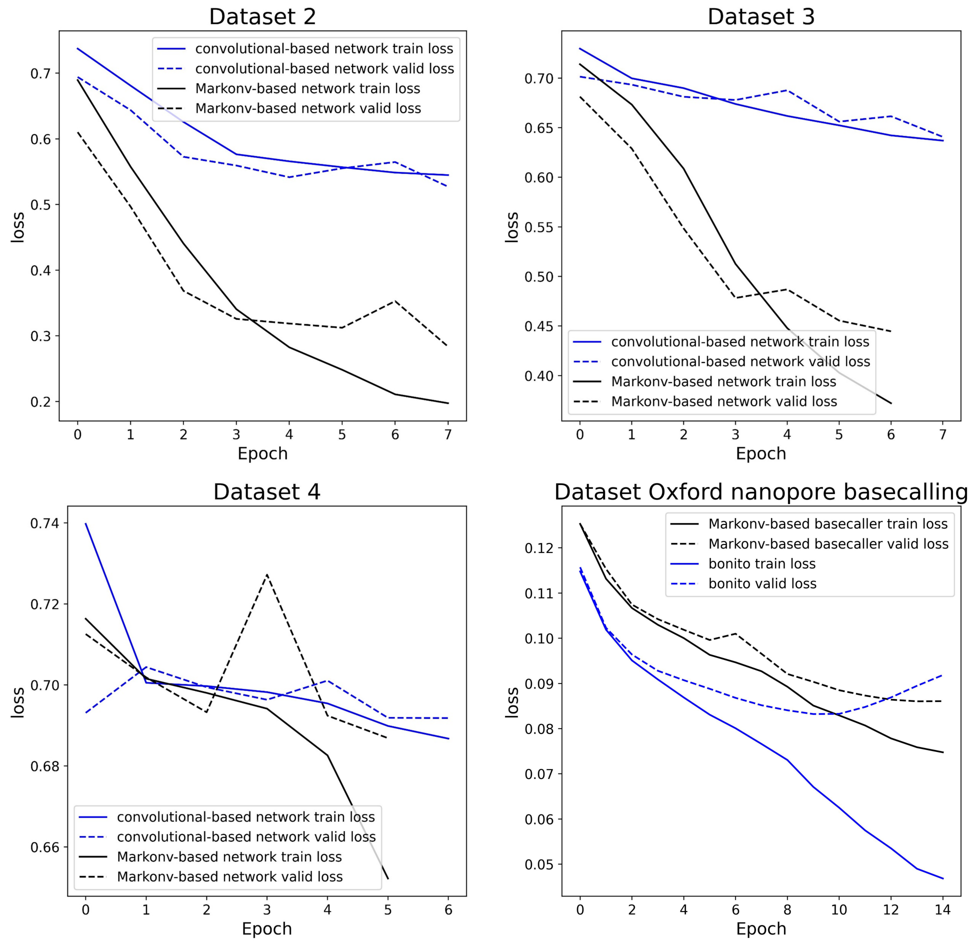


Figure 6: The convergency of Markonv-based networks (for simulation datasets) / the Markov-based basecaller (for Oxford Nanopore sequencing datasets) and convolutional-based networks (for simulation datasets) / the Bonito model (for Oxford Nanopore sequencing datasets). The x-axis is the number of epochs, while the y-axis is the loss for the training or validation dataset.
